# Robust learning-driven structural and functional plasticity of spines in the mature mouse cortex

**DOI:** 10.64898/2026.05.19.726233

**Authors:** Mona Fariborzi, Delaney G. Eaves, Lara Y. Demir, Yuxi Wu, Adam Skeens, Martin Hruska, Adema Ribic

## Abstract

Spines in the adult cortex are thought to be highly stable, and that their capacity for modest remodeling supports learning. Using a visual association task and a multilevel imaging approach in adult mice, we found a robust learning-driven increase in the complexity of spine nanostructure, as well as a rapid and persistent increase in spine formation during task acquisition that were accompanied by an overall reduction in spine size of layer 2/3 neurons in the primary visual cortex (V1). Trained animals further had an increased fraction of spines tuned to the task-relevant orientations, and the discriminability of spine responses in naïve mice was predictive of their subsequent performance. Our results demonstrate that learning drives an increase in spine preferences for task-relevant information and point to reconfiguration of spine nanostructure and spine inputs as the structural drivers of these changes.

## Introduction

Excitatory, glutamatergic inputs in mammalian cortices predominantly form connections (synapses) with spines, membranous postsynaptic compartments that protrude from the dendrites (1–3). Cortical spines display substantial structural plasticity and high turnover rates postnatally, contrasted by heightened stability in adulthood (4–7). However, previous studies in adult mice demonstrated that learning new skills or associations results in increased spine formation, turnover, and structural plasticity. For example, on apical tufts of layer 5 neurons, fear conditioning results in new spine formation in the auditory cortex (8), and increased clustering in the retrosplenial cortex (9). Similarly, in the motor cortex layer 5 neurons and in the visual cortex layer 2/3 neurons, rotarod training and visual perceptual learning, respectively, result in increased spine formation (10,11). Additionally, in both layer 2/3 and layer 5 neurons of the barrel cortex, learning a whisker detection task results in increased spine density, and increased spine head diameter in layer 5 neurons specifically (12). These studies hence suggest that increases in spine density, head size, and clustering support learning, but it is unclear how these structural changes functionally relate to distinct components of newly learned skills or associations.

Spines can be studied functionally using calcium indicators or glutamate sensors, such as gCaMPs or iGluSnFR respectively (13–16). Studies using such sensors reveal the functional changes in spines associated with learning. For example, mice learning a motor task have increased activity of newly formed and existing spines in the primary motor cortex (M1), as well as increased correlations of spine activity with movements during task performance (15,17). However, the reported changes in spine activity are not associated with specific motor components of the task (17). While this finding may simply reflect the difficulty of sampling a sufficient number of spines to capture the complex functional organization of M1 (18), the link between learning-related spine formation and specific task components remains unclear.

In the primary sensory areas, neurons represent a defined feature in the sensory space. In the primary visual cortex (V1), layer 2/3 neurons and their spines are selective for oriented edges (13,19), and visual association tasks that rely on orientation discrimination result in increased representation of task-relevant orientations in V1 neurons (20–23). Hence, if functional changes in spines during learning indeed support task performance, the representation of the task-relevant orientations should be reflected in spine orientation preferences as well. To address this, and test if learning-driven changes in spine structure and function are associated with specific components of a task, we investigated learning-driven structural and functional plasticity of spines on layer 2/3 neurons in the mouse V1. Using confocal, super-resolution, and *in vivo* structural and functional imaging of fluorescently labeled spines, we found that learning increased spine formation, the number of putative synapses onto spines, and their preference for task orientations, providing further evidence for the causal involvement of spines in learning.

## Results

As structural plasticity is a hallmark of learning, we first sought to identify structural changes in the spines of the visual cortex associated with learning. To do so, we injected newborn (postnatal day 0) mice intraventricularly with diluted AAV-hSyn-GFP to achieve sparse transduction (24) and waited until the animals were 2 months old to start the training. We then randomly assigned them to naïve or trained groups. For the trained group, water-restricted, head-fixed mice were trained on a visual associative task in which they learned to associate a drifting grating (120°) orientation with a water reward, while a dissimilar orientation (60°) was not paired with anything (Fig 1A). Mice were trained every day for two 30-minute, 100 trial sessions. The naive animals underwent identical headpost surgery, water-restriction, and exposure to the training environment where they received their daily allotment of water but were not exposed to any visual stimuli.

**Fig 1.**
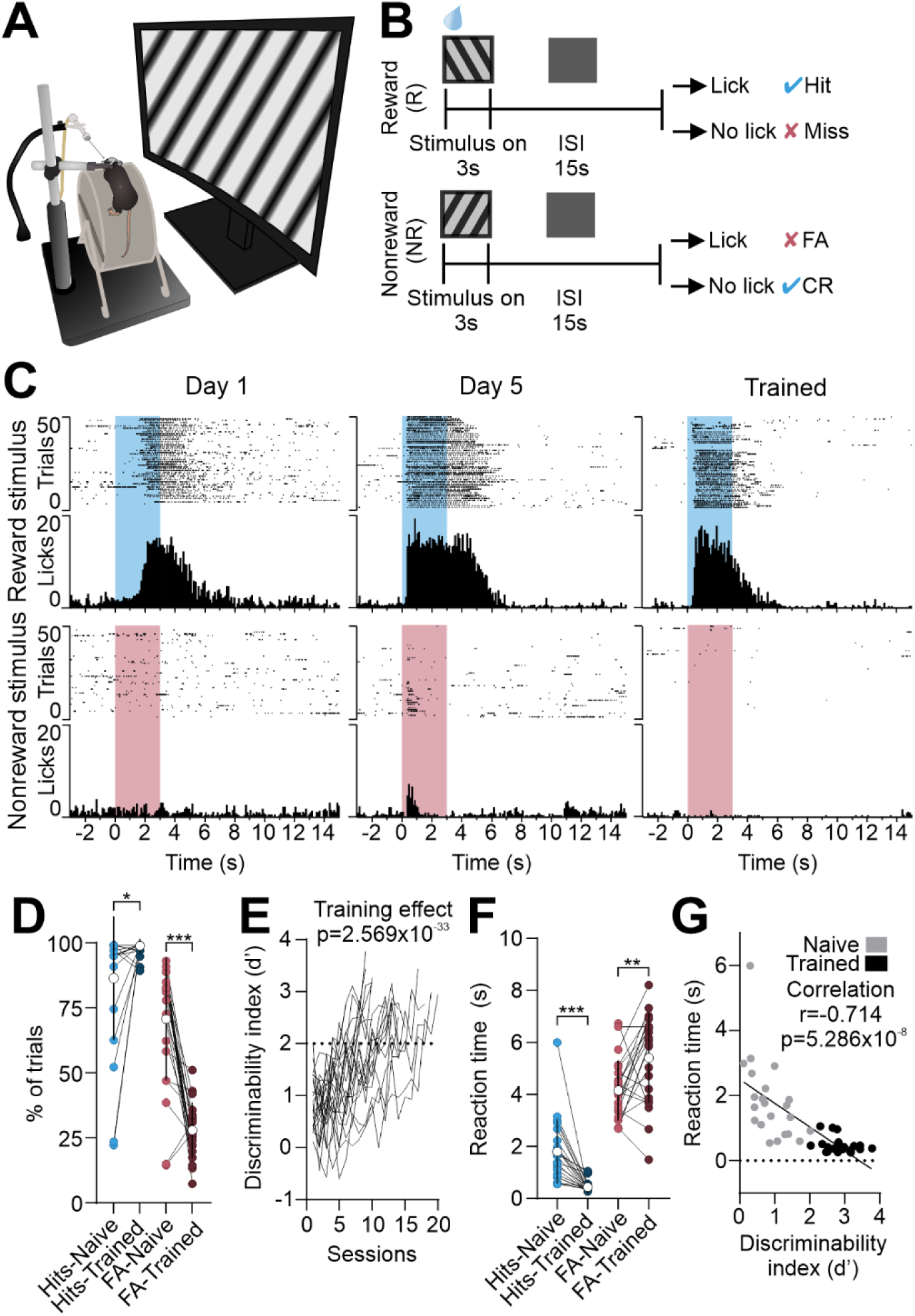
Adult mice learn to differentiate between visual cues. **A.** Task schematic of head-fixed adult mouse with stimulus displayed on the computer monitor centered on the right eye. **B.** Trial structure schematic. The visual stimulus is on for 3 seconds followed by a 15 second interstimulus interval (ISI). The rewarded stimulus is an orientation grating of 120°, and non-rewarded stimulus is 60°, while the screen is blank during the interstimulus interval. Water is dispensed at the onset of the rewarded visual stimulus. Four outcomes to this task are depicted on the right. **C.** Example lick raster and peristimulus time histograms (PSTHs) for a single mouse across 3 different training timepoints: first training session, day 5 of training, and final training session. The colored bars show when the visual stimulus is on, with the blue bars representing the rewarded stimulus, and red representing the non-rewarded stimulus. **D.** The percentage of hit trials for all the rewarded trials (blue), and the percentage of false alarm trials for all the non-rewarded trials (red) during the naive and trained session. Each dot represents an individual animal (Repeated measures ANOVA, F(^1,21^)= F_training_=7.882, F_response_=935.732, F_training*response_=228.02. Post hoc: hits p=0.027, FA p=3.334×10^-6^, N=22 animals). **E.** The discriminability index (d’) across training sessions. Each line is the d’ for an individual mouse (Linear mixed model (LMM), Training effect p=2.569×10^-33^, N=22 animals) **F.** The time to first lick (reaction time) for hit (blue) and false alarm (red) trials during the first and last training sessions (Repeated measures ANOVA, F(^1,21^)= F_training_=0.041, F_response_=180.589, F_training*response_=48.219. Post hoc: hits RT p= 9.187×10^-5^, FA RT p=0.004, N=22 animals). **G.** The correlation between d’ and reaction time to reward trials for naive and trained mice. Black dots represent naive animals, and gray dots represent trained animals (Pearson’s correlation, r=-0.714, p=5.286×10^-8^, N=22 animals). *p<0.05,**p<0.01,***p<0.001

For quantification of performance, we used the classical signal detection theory(25), where the task has 4 outcomes: hit (licking to the rewarded stimulus), miss (not licking to the rewarded stimulus), false alarm (FA, licking to the non-rewarded stimulus), and correct rejection (CR, not licking to the non-rewarded stimulus, Fig 1B). Across training days, mice became increasingly responsive to the rewarded stimulus and learned to withhold licks to the non-rewarded stimulus (Fig 1C). This was reflected in the number of hit trials significantly increasing and false alarm trials significantly decreasing in trained mice (Fig 1D, Repeated measures ANOVA, F(^1,21^)= F_training_=7.882, F_response_=935.732, F_training*response_=228.02. Holm post hoc: hits p=0.027, FA p=3.334×10^-6^, N=22 mice). The increase in performance was measured as discriminability index (d’)=z(H)-z(FA), which progressively increased during training (Fig 1E, Linear mixed model (LMM), Training effect p=2.569×10^-33^, N=22 animals). The animals were considered fully trained once they achieved a d’ of 2 or above for 3 sessions in a row (Fig 1E), which on average was achieved after about 12 sessions (SD=±3.9 sessions, 22 mice). In our task, recognizing the nonreward cue is important for increased task performance, illustrated by a negative correlation between the fraction of false alarms and d’ (S1 Fig). Increase in performance was also reflected in the reaction times, which significantly decreased for hits and increased for FA trials in trained mice (Fig 1F, Repeated measures ANOVA, F(^1,21^)= F_training_=0.041, F_response_=180.589, F_training*response_=48.219. Holm post hoc: hits RT p= 9.187×10^-5^, FA RT p=0.004, N=22 mice). Further, the reaction times to the rewarded stimulus were correlated with task performance (Fig 1G, Pearson’s correlation, r=-0.714, p=5.286×10^-8^).

After training or the matched amount of time for naïve mice, animals were perfused, and their brains isolated and sectioned. Using confocal microscopy, superficial layer 2/3 neurons in the visual cortex were identified based on distance of their cell body from the cortical surface (∼150-200 microns deep) and spines on the tertiary branches of the basal or apical dendrites were imaged (Fig 2A-B). First, spine density was quantified for the basal and apical branches, and no significant difference was observed between naive and trained animals, though there were overall fewer spines on the basal branches (Fig 2C, S2 Fig A, LMM, Branch location p=0.004, Training p=0.924, N=5 naive, 7 trained mice). As the spine head size can also change in an experience-dependent manner (12,15,17,26), we next measured the head diameters of identified spines and found that there was a higher frequency of spines with a smaller head size for both apical and basal branches (Fig 2D, Kolmogorov-Smirnov test, Apical p=2.2×10^-16^, D=0.2162, Basal p=2.2×10^-16^, D=0.19649, N=1564 spines naive apical, 2153 spines trained apical, 1394 spines naive basal, 1759 spines trained basal).

**Fig 2.**
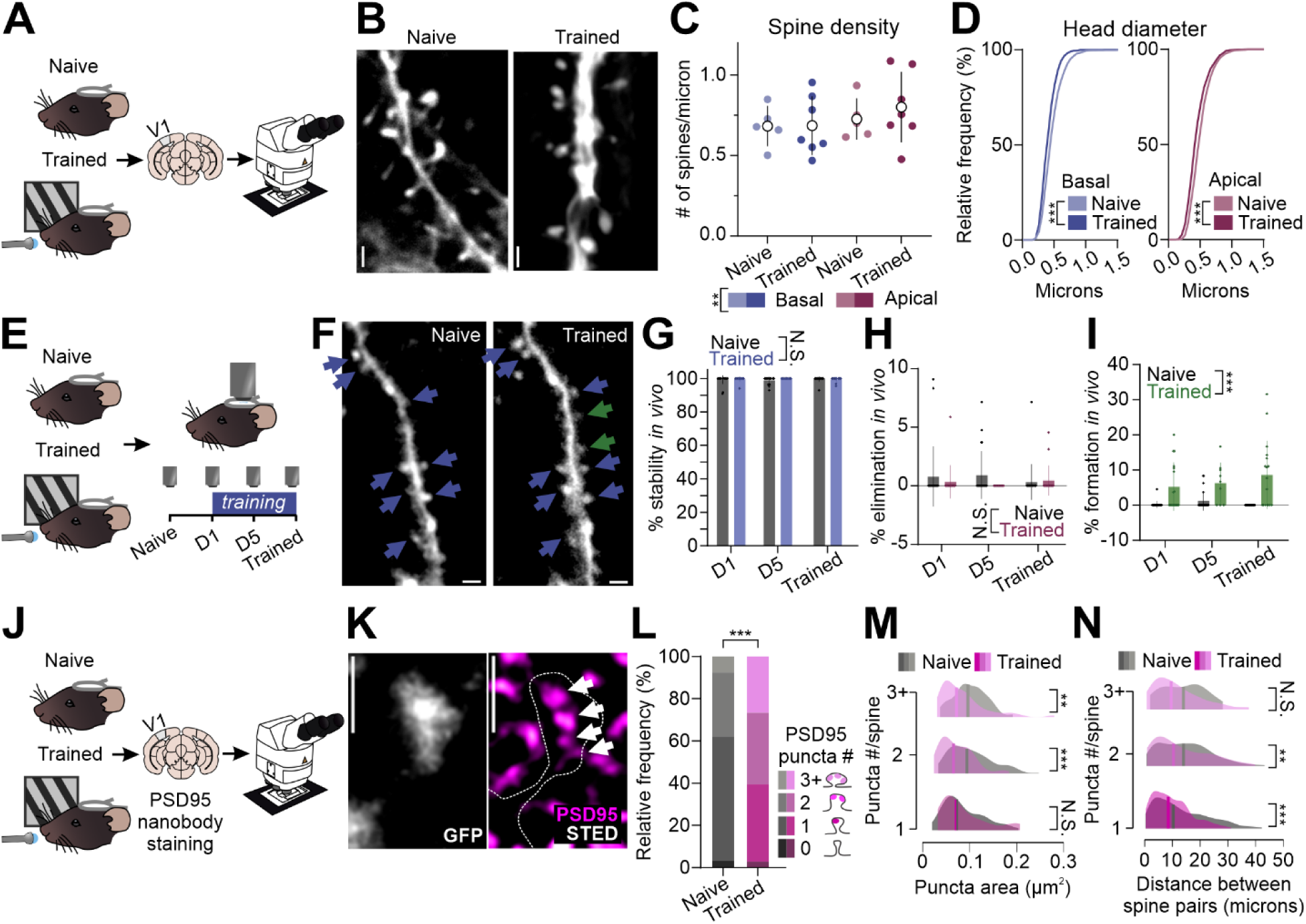
Learning results in changes in head diameter, spine formation, and density of PSD95 puncta per spine. **A.** Schematic of the confocal experiment. Mice were separated into naive or trained groups, and after the trained group learned the task, both groups were perfused, and brains were sectioned. Spines of layer 2/3 neurons of the V1 were imaged using confocal microscopy. **B.** Representative image of a dendritic branch of layer 2/3 neuron in naive (left) and trained (right) mice. Scale bar: 1 micrometer. **C.** Spine density quantification for naive and trained group for basal (blue) versus apical (magenta) branches (LMM, Branch location p=0.004, Training p=0.924, N=5 naive, 7 trained mice). **D.** Cumulative frequency histograms of head diameters for basal (blue) and apical (magenta) branches (Kolmogorov-Smirnov test, Apical p=2.2×10^-16^, D=0.2162, Basal p=2.2×10^-16^, D=0.19649, N=5 naive, 7 trained mice). **E.** Schematic of *in vivo* 2-photon imaging of GFP labeled branches of layer 2/3 neurons in V1. Mice were separated into naive and trained groups, and the same dendrites were imaged before the animal received any training, after the first training session, after 5 days of training, and once the animal was fully trained. The naive animals were imaged during the same time points. **F.** Representative image showing the same branch imaged during the naive and trained timepoints. Blue arrows show stable spines, and green arrows show newly formed spines. Scale bar: 1 micrometer. **G-I.** The percentage of stable (G), eliminated (H), and newly formed (I) spines in the naive group (gray) versus trained group (color) across the imaging time points ((G) LMM, Training effect p=0.279, (H) LMM, Training effect p=0.338, (I) LMM, Training effect p=2.347×10^-4^, N=6 naive, 6 trained mice). **J.** Schematic of the STED experiment. Mice were once again separated into naive or trained groups and once the trained animals learned the task, both groups were perfused, V1 sections isolated and stained using PSD95 nanobodies. The dendrites of layer 2/3 neurons were imaged using STED microscopy to quantify both the spines, as well as the PSD95 nanoclusters. **K.** Representative image showing a spine (gray, left) and the PSD95 puncta (magenta) within the spine (dotted line, right). White arrows point to PSD95 puncta. Scale bar=1 micrometer. **L.** Relative frequency of spines with 0,1,2, or 3+ PSD95 puncta in the naive (grey) or trained (magenta) group (Chi-Square test, p= 0.0008, N=187 spines naive, 139 spines trained, 3 mice per group). **M.** Distribution of puncta area between naive (gray) or trained (magenta) groups for spines with 1,2, or 3+ puncta (Kolmogorov-Smirnov Test, 1 puncta D=0.15488, p=0.3003, 2 puncta D=0.28331, p=0.0005119, 3 puncta D=0.33146, p=0.001718). **N.** The difference in distance between spines with 1, 2 or 3+ PSD95 puncta between the naive (gray) or trained (magenta) group (Kolmogorov-Smirnov test, Spines with 1 puncta p=6.624×10^-8^, D=0.185, Spines with 2 puncta p=0.007768, D=0.1463, Spines with 3+ puncta p=0.2553, D=0.22807). *p<0.05,**p<0.01,***p<0.001

As several previous studies reported an increase in spine density after learning in layer 5 neurons using Thy1-GFP mouse line (8–10,27), we repeated the imaging of structural changes in our task paradigm in Thy-GFP mouse line. We found that training had an overall significant effect on spine density, though it was not significantly different when specifically comparing apical or basal branches (S2 Fig B, S3 Fig A, LMM, Training effect p=0.047, AIC=-55.751, BIC=-24.892, Post hoc Apical Naive vs Trained p=0.127, Basal Naive vs Trained p=0.127, N=7 naive, 7 trained mice). Furthermore, we found that the head diameter of layer 5 spines was significantly increased on both basal and apical branches in trained mice (S3 Fig B, Kolmogorov-Smirnov test, Apical p=2.2×10^-16^, D=0.19603, Basal p=2.2×10^-16^, D=0.25847, N=2589 spines naive apical, 2985 spines trained apical, 2170 spines naive basal, 2381 spines trained basal). Altogether, our results from confocal imaging indicate layer-specific structural changes after visual associative learning in adult V1, with an overall reduced spine head size in layer 2/3 neurons and an increase in spine density and head size in layer 5 neurons.

Baseline structural plasticity of spines in the adult brain is low, but present and measurable on the scale of days when spines are imaged longitudinally *in vivo* using 2-photon imaging (28). As such changes cannot be captured *ex vivo* (as in Fig 2C-D), we next performed *in vivo* 2-photon longitudinal imaging of spines across training to quantify the impact of learning on structural plasticity of spines on individual, longitudinally tracked dendritic branches. We chose to focus on layer 2/3 as our *ex vivo* results indicated that learning had little impact on the structure of its spines, despite layer 2/3 neurons receiving both feedforward inputs from layer 4 of V1, as well as long-range inputs that facilitate the acquisition of visual associative tasks (29–31). Furthermore, neurons in layer 2/3 of V1 have a higher level of orientation selectivity compared to layer 5 (32). We once again injected newborn mice intraventricularly with AAV-hSyn-GFP and once they reached 2 months of age, a cranial window surgery was performed in which the skull above the V1 was removed and a glass window was secured in its place with a headpost. Mice were once again separated into either the naive or trained group, and branches from layer 2/3 neurons were identified and imaged before exposure to the training environment (naïve) or training (trained). In trained mice, the same branches were tracked after the first training session, after the fifth training day (training mid-point), and after the animal fully learned the task (Fig 2E, F). Naïve mice followed the same imaging schedule: day 1, 5 and last day of training for the trained mice of that cohort.

The stability of spines identified in the first timepoint was not significantly different between naive and trained animals and remained very high throughout all timepoints (Fig 2G, LMM, Training effect p=0.279, N=6 naive, 6 trained mice). Spine elimination was also very minimal and was not significantly different between the groups for any of the timepoints (Fig 2H, LMM, Training effect p=0.338, N=6 naive, 6 trained mice), in agreement with high stability of spines in the visual cortex of adult mice (5,6,28). However, in trained mice, just one training session significantly increased new spine formation, which remained high throughout the training with 76% of branches gaining new spines (Fig 2I, LMM, Training effect p=2.347×10^-4^, N=6 naive, 6 trained mice). Our longitudinal 2-photon imaging hence demonstrates that learning induces new spine formation in layer 2/3 neurons of the V1, overall, population-level spine density being maintained.

As our confocal imaging found changes in the spine head size between naïve and trained mice, we next sought to determine whether the nano-architecture of individual spines is altered after learning. Previous work demonstrated that structural plasticity is linked to changes in the molecular nano-organization of dendritic spines, where increase in the size of dendritic spines was associated with the increase in numbers, but not size, of individual PSD-95 nano clusters (33,34). To do this, we injected adult mice with AAV-cAMKII-GFP in layer 2/3 of the visual cortex, implanted the headposts and randomized mice into the naive or trained groups. After training (or exposure to the training environment, as for Fig 2A), we perfused and sectioned their brains to 10 µm thickness (Fig 2J). Because of the importance of PSD-95 nano-organization for structural plasticity (33,34), the brain sections were probed with nanobodies against PSD-95 to label the postsynaptic nano-domains of the synapse (35). We then performed super-resolution imaging using 3D-STimulated Emission Depletion (STED) microscopy to resolve PSD95 nanoclusters within spines (33,35). For each identified spine, we quantified the number of PSD95 puncta that was found in the same plane for the naive and trained groups and found that trained animals have a significantly higher frequency of spines with multiple PSD95 puncta (Fig 2L, Chi-Square test, p= 0.0008, N=187 spines naive, 139 spines trained, 3 mice per group). We also quantified the head diameter of spines in relation to the number of PSD95 puncta. Consistent with previous work (33,34,36,37), we found a positive correlation between spine head size and the number of PSD95 puncta (S4 Fig A, Pearson’s correlations, Naive Pearson’s r= 0.5516, p=5.55×10^-20^, Trained Pearson’s r= 0.5348, p=3.350×10^-16^). Though both the naive and trained groups had a positive correlation, there was a significant reduction in the elevation but not slope of the linear regression line between the naive and trained groups, indicating that trained mice has a smaller size of PSD95 nanodomains (S4 Fig A, Simple linear regression, Slope F=1.892, DFn=1, DFd=322, p=0.1699, Elevation F=6.071, DFn=1, DFd=323, p=0.0143). To further investigate this difference, we compared the distribution of puncta area on spines with 1, 2 or 3+ (3 and 4) nanodomains. Here, we found a significant shift in the distributions of puncta area for spines with 2 and 3+ puncta towards smaller values, but not spines with 1 nanodomain (Fig 2M, Kolmogorov-Smirnov Test, 1 puncta D=0.15488, p=0.3003, 2 puncta D=0.28331, p=0.0005119, 3 puncta D=0.33146, p=0.001718). Furthermore, there was a significant effect of training that was dependent on the number of puncta per spine, with an area reduction of spines with 2 puncta (S4 Fig B, LMM, Training effect p=0.408, Puncta # effect p=0.99, Training*Puncta # effect p=0.047, Naive 2 puncta spine vs Trained 2 puncta spine P_holm_=6.876×10^-4^ N=187 spines from 3 naive mice, 139 spines from 3 trained mice). We also quantified the distance between spines with 1, 2, or 3+ PSD95 puncta, as spines tend to cluster together after learning. We found that spines with 1 and 2 PSD95 puncta were more clustered in trained animals, but not spines with 3+ puncta (Fig 2N, Kolmogorov-Smirnov test, Spines with 1 puncta p=6.624×10^-8^, D=0.185, Spines with 2 puncta p=0.007768, D=0.1463, Spines with 3+ puncta p=0.2553, D=0.22807). Our results hence demonstrate that learning leads to an increased nanoscale complexity of dendritic spines in the L2/3 pyramidal neurons despite reduced sizes of spines and their postsynaptic nanodomains.

Functionally, spine activity reflects the activity of the input it receives(16). Thus, we next asked whether the observed changes in spine nano-architecture would be reflected in altered spine activity after learning. To test this, adult mice were injected with a 1:1 mixture of diluted (1:40,000) AAV-hSyn-Cre and gCAMP8s to achieve sparse expression (13). At the same time, we secured a cranial window above the V1 with a headpost, and mice underwent water-restriction after the recovery as described previously (38,39). Mice were first imaged before receiving any training. During the imaging session, mice were lightly anesthetized with isoflurane to help stabilize the imaging window. Cell bodies and segments of the dendritic branches of layer 2/3 neurons in the V1 were imaged while the mouse was viewing drifting gratings. 3-5 days after the initial imaging session, the mice began the training and once they were fully trained, the same branches were imaged again while the animal was passively viewing drifting gratings (Fig 3A-C). We first calculated the orientation selectivity (OS) preferences for all the neurons imaged and found that their OS changed between naïve and trained stages (Fig 3D). Furthermore, 7/10 neurons were selective in the first imaging session (prior to training onset), while only 4/10 neurons were selective after training (Fig 3D). We then calculated the OS preferences for the spines that were longitudinally tracked during the training and found that the fraction of spines tuned to the non-rewarded orientation decreased while the fraction of spines tuned to the rewarded orientation increased (Fig 3E, Kuiper test, p=0.05, N=49 spines in naïve timepoint, 40 spines in trained timepoint from 6 animals). We also calculated the OS preferences for spines that were non-longitudinally tracked, i.e. imaged in one time point but not the other. The same trend held in this population of spines, with a decrease in the fraction of spines tuned to the non-rewarded orientation and an increase in the fraction of spines tuned to the rewarded orientation (Fig 3F, Kuiper test, p=0.002, N=120 spines in naïve timepoint, 136 spines in trained timepoint from 6 animals). To confirm that our results were not due to poor fits obtained with the classical Gaussian fit of OS preference, we analyzed our data with the vector sum calculations of OS for the naive timepoint (40). We found that the results from both methods were positively correlated with each other (S5 Fig A-B, A Pearson’s correlation, r=0.715, p=1.114×10^-25^, N=156 spines, B, Pearson’s correlation, r=0.993, p=8.765×10^-6^, N=7 neurons). OSI and gOSI were also significantly positively correlated for spines, though for neurons, OSI calculations overrepresented an OSI of 1 which has been previously reported (40)(S5 Fig C-D, C, Pearson’s correlation, r=0.573, p=1.039×10^-15^, N=156 spines, D, Pearson’s correlation, r=-0.082, p=0.861, N=7 neurons).

**Fig 3:**
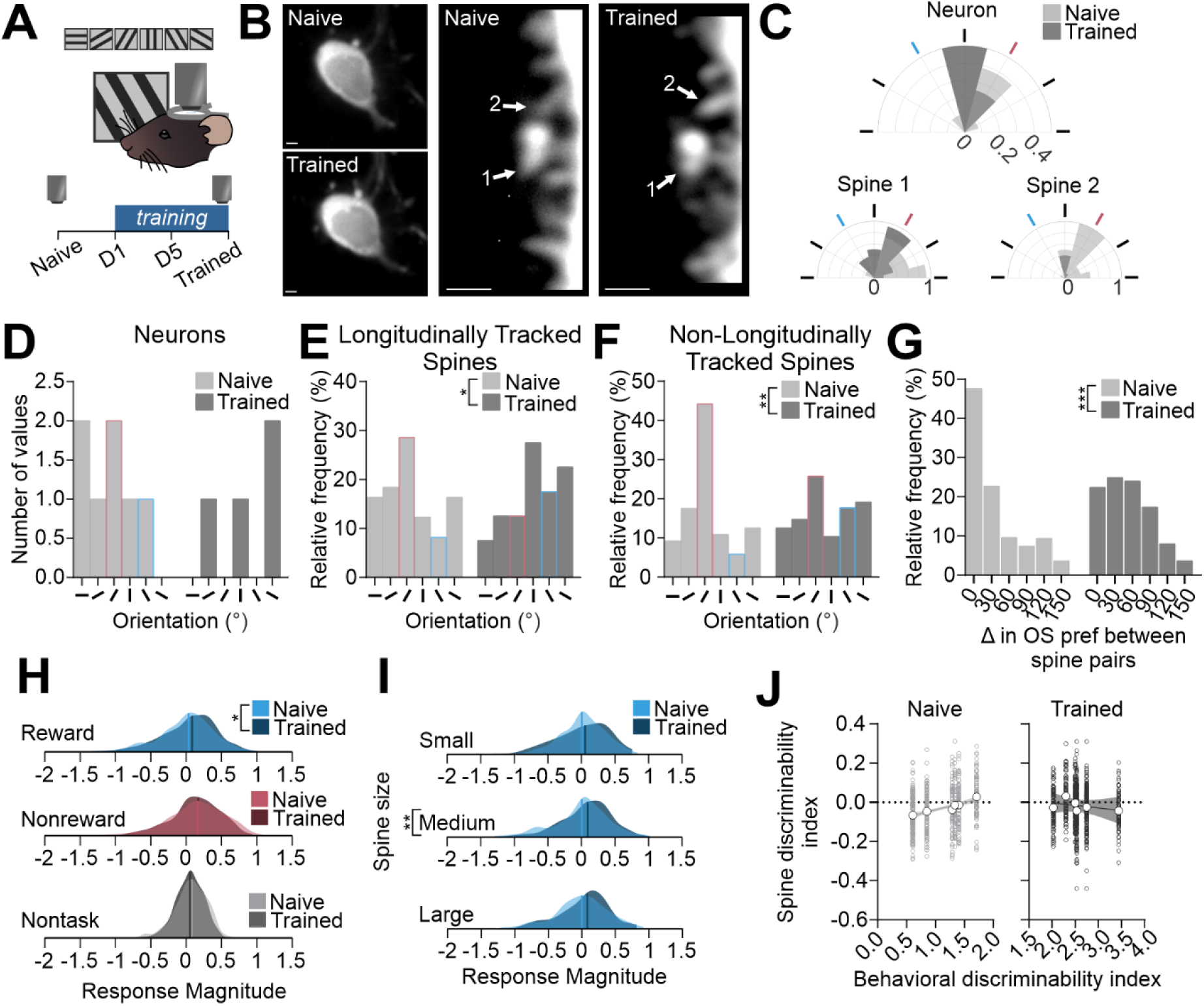
Learning results in increased representation of the rewarded cue in spines of layer 2/3 neurons in the V1. **A.** Schematic of experiment. Neurons sparsely labeled with gCaMP8s in the V1 were imaged while the mouse (under anesthesia) viewed drifting gratings of 6 orientations. 3-5 days after the initial imaging session, training began and once animals learned the task, the same dendrites were imaged. **B.** Representative images (standard deviation projection) of the same neuron and branch during the naive and trained imaging session. Scale bar: 1 micrometer. **C.** Polar plots for the neurons and spines in B. The rewarded orientation (120) is colored blue and the nonrewarded orientation (60) is colored red. **D.** Frequency distribution of preferred orientations of selective neurons in the naive (light grey) and trained (dark grey) group. Rewarded and nonrewarded orientations are denoted by blue and red outlines, respectively (N=10 total neurons from 6 mice, 7 selective neurons in naive timepoints, 4 selective neurons in trained timepoint). **E.** Frequency distribution of preferred orientations of longitudinally tracked spines in the naive (light grey) and trained (dark grey) group. Rewarded and nonrewarded orientations were denoted by blue and red outlines, respectively (Kuiper test, p=0.05, N=49 spines in naïve timepoint, 40 spines in trained timepoint from 6 animals). **F.** Frequency distribution of preferred orientations of non-longitudinally tracked spines in the naive (light grey) and trained (dark grey) group. Rewarded and nonrewarded orientations were denoted by blue and red outlines, respectively (Kuiper test, p=0.002, N=120 spines in naïve timepoint, 136 spines in trained timepoint from 6 animals). **G.** Frequency distribution of differences in orientation for spine pairs on a given branch during the naive and trained groups (Kolmogorov-Smirnov Test, p=2.2×10^-16^, D=0.28499, N=710 spine pairs naive, 555 spine pairs trained). **H.** Frequency distribution of response magnitude of spines to the rewarded, non-rewarded, and non-task orientations during the naive and trained imaging session (Kolmogorov-Smirnov, Rewarded orientation p=0.01781, D=0.080499, Non-rewarded orientation p=0.1947, D=0.056535, Non-task orientations p=0.08895, D=0.065364, N=648 spines Naive, 833 spines Trained). **I.** Frequency distribution for the response magnitude of spines to the rewarded orientation binned by small, medium, or large spines during the naive and trained imaging session (Kolmogorov-Sminorv Test, small group D=0.091322, p=0.4389, medium group D=0.11761, p=0.00682, large D=0.13445, p=0.2739, N=648 spines Naive, 833 spines Trained) **J.** Correlation between behavioral discriminability index and spine discriminability index during the naive (left panel, light gray) and trained (right panel, dark gray) imaging session (Pearson’s correlation, Naive r=0.278, p= 7.875×10^-13^, N=648 spines, 6 animals, Trained r=-0.076, p=0.028, N= 833 spines, 6 animals). *p<0.05,**p<0.01,***p<0.001.

In naïve animals, most orientation selective spines on V1 neurons have similar tuning preferences, in which they prefer the same orientation as the host neuron (13). As our results demonstrated an increase in the fraction of spines tuned to the task-relevant orientations after training, we hypothesized that spines in trained animals would have less similar orientation preference. To test this, we quantified the difference in orientation preferences for every spine pair on a given branch before and after training. During the naive imaging session, a substantial majority of spine pairs had similar orientation preferences, in agreement with previous reports (13,41) (Fig 3G). After training, however, the number of spines tuned to the same orientation significantly decreases, and the variability in orientation preferences increases, in agreement with our hypothesis (Fig 3G, Kolmogorov-Smirnov Test, p=2.2×10^-16^, D=0.28499, N=710 spine pairs naive, 555 spine pairs trained). Furthermore, we confirmed that the largest fraction of spines was tuned to the same orientation as the host neuron’s preferred orientation in both naïve and trained animals (S6 Fig)(13), indicating that the changes in the representation of task-relevant orientations at the level of neuronal population are due to changes in their inputs(42). The overall number of spines that display orientation selectivity was in line with previous findings in naïve animals (13) and not different in trained animals regardless of whether the spine was longitudinally tracked or not (S7 Fig, All spines, LMM, p=0.602, N=23 branches; longitudinally-tracked spines: Repeated measures ANOVA, F(^1,14^)= F_training_=2.179, p=0.162, N=13 branches, non-longitudinally tracked spines: LMM, p=0.645, N=23 branches).

Formation of additional PSD-95 nanoclusters in individual spines could potentially alter the tuning curve of spines depending on the tuning preference of their inputs. Because we found a larger fraction of spines with multiple PSD-95 nanodomains using STED imaging (Fig 2L), we tested if training affected the orientation selectivity index (OSI) of spines. We found no significant differences in the OSI for either longitudinally or non-longitudinally tracked spines (S8 Fig A-D, Kolmogorov-Smirnov Test, longitudinally tracked spines, p=0.1739, D=0.20673, N=49 spines naive, 40 spines trained, non-longitudinally tracked spines, p=0.09933, D=0.15406, N=120 spines naive, 136 spines trained). The tuning width was also not significantly different between the naive and trained animals (S8 Fig E-F, Kolmogorov-Smirnov Test, longitudinally tracked spines, p=0.4306, D=0.17949, N=49 spines naive, 40 spines trained, non-longitudinally tracked spines, p=0.182, D=0.13763, N=120 spines naive, 136 spines trained).

Given the structural changes in head size of layer 2/3 spines after learning in the confocal data, we next wanted to investigate whether the same would be found in our functional imaging data. Consistent with the confocal imaging data (Fig 2D), there was a higher frequency of spines with smaller head diameters after learning (S9 Fig A, Kolmogorov-Smirnov Test, p=4.223×10^-5^, D=0.12153, N=648 spines naive, 833 spines trained). Moreover, this increased frequency of spines with smaller head diameters after learning was specific to spines tuned to the rewarded and non-task orientations, as well as non-selective spines, but was not present for spines tuned to the nonrewarded orientation (S9 Fig B-E, Kolmogorov-Smirnov Test, reward tuned spines p=0.01288, D=0.49876, N=13 spines naive, 31 spines trained, non-reward tuned spines p=0.4304, D=0.16382, N=67 spines naive, 41 spines trained, non-task orientation tuned spines p=0.0115, D=0.22782, N=86 spines naive, 103 spines trained, nonselective spines p=1.33×10^-5^, D=0.1492, N=451 spines naive, 659 spines trained). The reduction in head size with no changes in spine density was consistent across the confocal, STED, and *in vivo* imaging experiments, with a significant effect between control and trained animals for size measurements, but not experimental methods (spine density: S10 Fig A, LMM, Training effect p=0.128, Experiment effect p=0.626, N=107 branches Naive confocal, 145 branches Trained confocal, 5 branches STED, 14 branches 2P *in vivo*; spine size: S10 Fig B, LMM, Training effect p=0.011, Experiment effect p=0.748, N= 2958 spines naive confocal, N=3909 spines trained confocal, Estimated marginal mean=0.453, N=187 spines naive STED, Estimated marginal mean=0.497, N=139 spines trained STED, Estimated marginal mean=0.461, N=612 spines naive 2P *in vivo,* Estimated marginal mean=0.503, N=833 spines trained 2P *in vivo,* Estimated marginal mean=0.468).

We next compared the response magnitude of the spines to the rewarded, non-rewarded, and orientations not used in the task for both timepoints. Stimulus had a significant impact on the magnitude (S11 Fig, Generalized linear mixed model, stimulus p=1.307×10^-5^, N=648 spines Naive, 833 spines Trained), with an increased magnitude of responses to the rewarded stimulus only in trained mice (Kolmogorov-Smirnov p=0.01781, D=0.080499), but no significant differences for the nonrewarded or non-task orientations (Fig 3H). Given that there was an increase in spines with stronger response magnitudes to the rewarded orientation after training, we were interested in whether there may be a relationship between spine size and response magnitude. The spines were split into three groups, spines with small (0.2-0.4 µm), medium (0.4-0.6 µm), and large (>0.6µm) head sizes. We found that the increase in response magnitude after training was specifically for spines in the medium size group (Fig 3I, Kolmogorov-Sminorv Test, small group D=0.091322, p=0.4389, medium group D=0.11761, p=0.00682, large D=0.13445, p=0.2739).

Finally, we were interested in whether the functional changes in spines were correlated with learning. We calculated a discriminability index value for each spine based on the previous work (21). This measure subtracts the averaged response of a spine to the non-rewarded stimulus from the response to the rewarded stimulus and divides it by the pooled standard deviation (21). A value above 0 means the spine is more responsive to the reward cue, and a value below 0 means the spine is more responsive to the nonrewarded cue. During the naive imaging session, mice with a higher spine discriminability index were correlated with having better task performance during the first training session (Fig 3J, Pearson’s correlation, r=0.278, p= 7.875×10^-13^, N=648 spines, 6 animals). Conversely, during the final training session when the animal is an expert in the task, a lower spine discriminability index is correlated with better behavioral performance but only when comparing spines and not animals (Fig 3J, Pearson’s correlation, r=-0.076, p=0.028, N= 833 spines, 6 animals). We found no significant correlations between structural changes and behavior (S12 Fig), suggesting that while the structure alone is not indicative of changes in behavior in adult mice (43), the functional changes are.

## Discussion

In this study, we investigated the structural and functional learning-driven plasticity of spines in the adult mouse visual cortex. Learning resulted in robust structural plasticity, including an increase in PSD95 nanodomains onto individual spines, as well as new spine formation. Functionally, there were more spines tuned to the rewarded task orientation and the rewarded orientation elicited responses of higher magnitude across all spines regardless of the preference, with an overall decrease in the similarity of tuning preferences on spines of the same branch (Fig 4).

**Fig 4:**
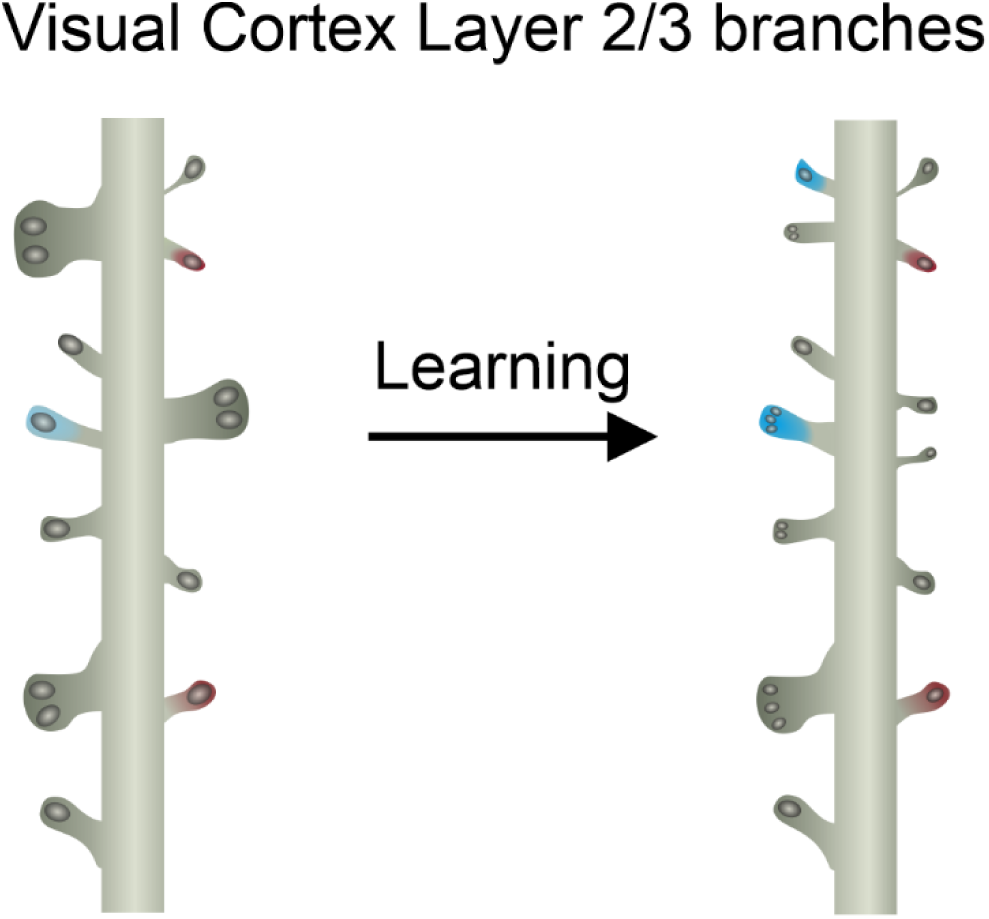
Learning-driven changes in V1 layer 2/3 spines. Given the overall structural stability of the adult brain, learning a visual task results in substantial functional changes, such as an increase in spine tuning preferences for the task-relevant orientations. While a large fraction of layer 2/3 neuron apical dendrites gains new spines, overall spine head sizes are reduced in trained animals, likely to accommodate the new spines. Parallel to the reduction in spine head size, the area of PSD95 nanodomains is also reduced in trained animals. However, trained animals have a larger fraction of spines with multiple PSD95 nanodomains, indicating that in adult animals, learning reconfigures synaptic nanostructure to facilitate the functional changes in spine response properties.

Spines in the adult brain have high stability compared to the developing brain (4,5,7,28). Furthermore, in the adult brain, different cortical areas have varying levels of spine turnover, with the visual cortex being highly stable compared to other areas (4,5). In line with these findings, our results also demonstrate heightened spine stability in adult V1, in both naïve and trained animals (44). Interestingly, we found an increase in new spine formation in our experimental paradigm, with no overall impact on spine density (Supplementary Fig 10A). While our results are in line with spine formation as a hallmark of learning (8,10,11), no change in overall spine density or spine loss in our data suggests that the increase in spine density might be branch- or neuron-specific, as has previously been previously shown (12,45,46).

Contrary to preserved stability of overall spine density, we found robust changes in spine structure and an increase in new spine formation on individual branches driven by learning. First, we found that training differentially impacts the head size of apical and basal branch spines on both layer 2/3 and 5 neurons in the V1, in line with previous reports (12). Second, our study found increased formation of new spines on layer 2/3 neurons in 76% of branches in trained animals. Finally, we found that spines in trained animals contain more postsynaptic apparatuses, identified through STED imaging of PSD95 nanoclusters. Spines are known to undergo substantial activity- and experience-dependent remodeling (47). In naive animals, super-resolution microscopy, such as STED and STORM, have been used to describe and quantify subsynaptic structures such as nanodomains and nanomodules. Nanodomains and nanocolumns represent subsynaptic specializations containing a higher density of transmitter receptors within the postsynaptic density and the trans-synaptic apparatus connecting it with the presynaptic release machinery, respectively (1,33,48,49). Our finding of increased frequency of spines with multiple post-synaptic densities (PSDs) in trained animals suggests that these spines contain more neurotransmitter receptors. Consistent with these findings, a previous study in the barrel cortex demonstrated that learning a whisker detection task results in increased surface expression of AMPA receptors in neurons activated during the task (12). While our super-resolution approach did not track the dynamics of the presynaptic compartments during learning, most PSDs in spines are opposed by a presynaptic marker, such as vGlut1 and vGlut2 (33,35). The increase in multi-PSD spines in trained animals hence indicates that the number of inputs onto single spines is robustly increased after learning, and future studies should aim to investigate the identity of these presynaptic inputs. Furthermore, our finding that there is increased clustering of spines with 1 and 2 PSDs suggests that spines may be forming new connections with already existing boutons. This hypothesis is further supported by the finding of smaller PSD areas for spines containing 2 and 3 PSDs in trained mice. Indeed, a previous study demonstrated that learning in adulthood increases synaptogenesis primarily through the addition of new inputs onto existing spines or through pairing newly formed spines with existing inputs (50,51).

Our findings of increased spine formation, a higher frequency of spines with a smaller head diameter, and in increase of spines with multiple, but smaller PSD areas in trained animals, suggests that new spine formation in layer 2/3 neurons is facilitated by the reductions in spine head size of existing spines, without impacting their function. To the contrary, a higher number of spines with multiple PSD95 nanodomains and a higher response magnitude to the reward orientation in trained animals, suggests that learning induces functional strengthening of spines that may not be reflected in spine head size alone. While a linear relationship between spine size and synapse strength has been reported (52), synapses show a large variability of their estimated functional strength, especially in the medium-sized PSD area group, which is where we see the largest functional changes in our study. Hence, spine enlargement may not be necessary for functionally increasing synaptic strength during learning, but reconfiguration of synaptic nano-structure (such as formation of new PSDs) might. Future studies that can track the same dendritic segments with STED and 2P imaging of activity will be able to address this hypothesis.

In case of spines that show feature selectivity, such as those in the V1, adding new inputs would result in changes in their feature selectivity. In layer 2/3 of the visual cortex, neurons receive inputs from layer 4 of the V1 and long-range inputs from other cortical areas (29–31,53). Previous work investigating neuronal changes in selectivity to task cues has shown that learning results in neurons increasing activity in response to the task cues (20,21,23,42). A potential mechanism for increased responsiveness to task cues is that neurons are receiving more inputs that are tuned to the task features. Indeed, we found pronounced changes in feature selectivity of spines longitudinally tracked during training, with an overall increase in the fraction of spines responding to the rewarded cue in trained animals. Furthermore, after training, spines had a larger response magnitude for the rewarded orientation but not the non-rewarded orientation or non-task orientations, suggesting that new inputs represent the learned information. Despite higher representation and response magnitudes to the rewarded stimulus, we found that increased spine discriminability to the nonrewarded stimulus was associated with better task performance in trained animals. Our results support that an increase in the representation of task cues in the V1 of trained animals is a result of learning-driven changes in neuronal inputs, and that these functional changes are necessary for learning as they correlate with performance in naïve and trained animals. The circuit origin of these inputs is unclear at this point. Our imaging sampled layer 2/3, which contains both local and long range, top-down inputs into the V1. As our approach used lightly anesthetized mice, it is likely that the new inputs into the V1 of trained animals are coming from another visual area given that the top-down areas do not show neuronal responses to the cues outside of task context (29,54,55).

Though learning resulted in differences in the distribution of orientation preferences of spines, the overall number of selective spines did not significantly change for either the longitudinally or non-longitudinally tracked spines (Supplementary Fig 7). Furthermore, we had a comparable number of selective spines found on the layer 2/3 neurons as were found in previous work, though different generations of gCaMP sensors were used (13). In line with this previous work, we also found that in the naive group, though there is a variety of tuning preferences for spines, most spines are tuned to the same orientation as the neuron. (Fig 3I, Supplementary Fig 6, (13)). Because our data had two neurons tuned to the 60° orientation in the naive group, we also found an overrepresentation of spines tuned to the 60° orientation in the naive group for both longitudinally and non-longitudinally tracked spines (Fig 3, D-F). However, we found that after learning, mice had more non-selective neurons, and a larger variation of tuning preferences amongst their spines (Fig 3I), which is in line with previous work showing that even non orientation selective neurons contained selective inputs (56). Interestingly, our finding of a reduced number of neurons responding to task orientations matches a previous report, despite a small number of neurons in our study (42).

Altogether, our study comprehensively investigated learning-driven plasticity of spines in the adult cortex at substructural, structural, and functional levels. We found structural plasticity in the form of new spine formation and a higher frequency of spines with multiple putative synapses. Functionally, we found that learning increased the representation of the rewarded stimulus amongst spines and increased the variability of tuning preferences amongst inputs. Future studies could expand these findings by identifying specific neurons that were active during the task and combining *in vivo* with *ex vivo* super-resolution techniques to link the structural and functional plasticity of spines on task-active neurons and correlate them with discrimination performance (12,15,17).

## Materials and Methods

### Mice

Mice were maintained on C57BL/6 and Thy1-EGFP background (The Jackson Laboratory, Bar Harbor, ME) on a reverse 12:12 light:dark cycle (lights off 11 AM-11 PM), with food and water *ad libitum,* except during visual discrimination training (see below). Animals of both sexes were used between 2 and 7 months of age. Animals were treated in accordance with the University of Virginia Institutional Animal Care and Use Committee guidelines.

### Viruses

Following viruses were purchased from Addgene: AAV-hSyn-eGFP (50465-AAV1; 5×10^12^ vg/ml), AAV-hSyn-Cre-P2A-dTomato (107738-AAV1; 1.3×10^13^ vg/ml), AAV-syn-FLEX-jGCaMP8s-WPRE (162377-AAV1; 2.1×10^13^ vg/ml), and AAV-CamKIIa-EGFP (50469-AAV9, 1×10^13^ vg/ml). All viruses were diluted in sterile PBS before the injection, and specific dilutions are indicated below.

### Surgeries

For intraventricular injections, mice were injected 6-12 hours after birth. Pups were taken from their dam and anesthetized with ice. NanoFil syringe (35 G beveled needle; WPI, Sarasota, FL) was used to inject the into the ventricle of the left hemisphere, which was located 2/5 of the distance from the lambda suture to the eye, and 3 mm deep. 2 µL of 1:10 diluted AAV-hSyn-eGFP was infused at a rate of 10 µL/min, as previously described (24). Pups were then placed on a heat pad to recover and returned to their dam. Animals were left in their home cage for at least 2 months, until adulthood.

For adult surgeries, animals were anesthetized with isoflurane in oxygen (2-2.5% induction, 1–1.5% maintenance), warmed with a heating pad at 38°C and given subcutaneous injections of Buprenorphine SR (1 mg/kg) or Rimadyl (5 mg/kg) and 0.25% Bupivacaine (beneath their scalp). Eyes were covered with Puralube (Decra, Northwich, UK). Scalp and fascia from Bregma to behind lambda were removed, and the skull was cleaned, dried and covered with a thin layer of Scotchbond adhesive (3M, Maplewood, MN). Skin edges were sealed with VetBond (3M).

For virus injections, the head was immobilized in a custom-made stereotactic apparatus and small craniotomies were made on the left hemisphere with a dental drill at the following coordinates: primary visual cortex (V1): 0.5 mm anterior to lambda, 2-3 mm lateral to midline. For cranial window surgery, a 3 mm circular craniotomy was made above the V1 coordinates. NanoFil syringe (35 G beveled needle; WPI, Sarasota, FL) was then used to inject the virus at 1 nl/s using a Syringe Pump (KD Scientific, Holliston, MA) at 2 different locations of the V1 with a depth of 100 µm beneath the brain surface for targeting layer 2/3 of the cortex. For the STED experiments, 50 nL of 1:10 diluted AAV-CamKIIa-EGFP was infused per injection site. For functional imaging, 150 nL of a 1:1 mixture of 1:40,000-1:100,000 AAV-hSyn-Cre-P2A-dTomato and undiluted AAV-syn-FLEX-jGCaMP8s-WPRE, respectively, was infused per injection site. The syringe was left in place for 5-10 minutes after the injection to allow the virus solution to diffuse and then very slowly raised to the surface to prevent the backflow. For the cranial window surgeries, a 5 mm circular glass coverslip was then placed over the craniotomy and secured in place with VetBond.

The head plate (stainless steel, SendCutSend, Reno, NV) was attached with dental cement (RelyX Ultimate, 3M) after the virus injection and treatment of the skull with Scotchbond (3M) adhesive. After the cement cured, the well of the head plate was filled with silicone elastomer (Reynold Advanced Materials, Brighton, MA) to protect the skull. If the mice received Rimadyl for analgesia, they were given Rimadyl dissolved in hydrogel (1% food-grade agar in distilled water) during the recovery. Animals were group-housed after the implantation and monitored daily for signs of shock or infection.

### Visual stimuli

For visual stimuli during the visual discrimination task, the blank screen was generated with MATLAB (MathWorks, Natick, MA) using the PsychToolBox extension (Brainard, 1997) and presented on a gamma corrected 27” LCD. The screen was centered 20-25 cm from the mouse’s right eye, covering ∼80° of visual space. Visual stimuli (orientations 120° and 60°) were presented at 0.15 cpd and 100% contrast in a random sequence for 3 seconds, followed by a 15 second long interstimulus interval (blank screen). Presentation of 120° triggered the delivery of 10 µl of water through a syringe pump (New Era Pump Systems, Farmingdale, NY), while the presentation of 60° had no consequence.

The visual stimuli for the functional imaging were presented on a gamma corrected 7” LCD screen. The screen was centered 20-25 cm from the mouse’s right eye, covering ∼80° of visual space. The screen was placed in a pyramidal enclosure made from light tight foil (ThorLabs, Newton, NJ) whose top opening was positioned over the mouse’s eye for the imaging sessions to reduce light contamination during the imaging. For calculating baseline orientation selectivity, 12 orientations 30° apart were presented at 100% contrast at 0.15 cpd, with 2 s long stimuli and 4 s interstimulus interval (blank screen). At least 10 repetitions of each orientation were shown.

### Visual Discrimination Task and Analysis

4-7 days after the headpost implantation, mice were gradually water restricted (from 3 ml of water/day to 1 ml of water/day) for 1 week. Food was available *ad libitum*. Mice were weighed daily and their weight was maintained at 85% of initial weight to prevent dehydration. During water restriction, mice were gradually habituated to handling by the experimenters, the treadmill, and the water delivery spout during daily habituation sessions. After 1 week of water restriction, the visual discrimination training began, where mice were head-fixed above the treadmill and a stainless-steel gavage needle (18G) was positioned near the mouse’s mouth for water delivery. The pump was controlled by the Power1401-3 data acquisition interface (CED) for lick detection. Mice were trained in 2 daily sessions consisting of 50 120° and 60° presentations each, for a total of 200 trials per day. Licks of the spout during the stimulus presentation, as well as during the first 10 s of the interstimulus interval were quantified according to the signal detection theory as Hits (the fraction of trials in which the mouse licked to the 120° orientation), Misses (the fraction of trials in which the mouse failed to lick to the 120° orientation), False Alarms (the fraction of trials in which the mouse licked to the 60° orientation) and Correct Rejections (the fraction of trials in which the mouse failed to lick to the 60° orientation), where the performance was measured as discriminability (d’)=z(H) -z(FA). Mice were trained until they achieved a d’ of 2 or above for 3 consecutive sessions.

### Histology, Imaging, and Analysis

For confocal imaging experiments, after the last training session, mice were anesthetized with Euthasol (pentobarbital sodium and phenytoin sodium) and transcardially perfused with warm 0.1□M phosphate buffer, followed by warm 4% paraformaldehyde (Electron Microscopy Sciences, Hatfield, PA). Brains were postfixed 1□hr at room temperature, followed by overnight fixation at 4□°C. Brains were sectioned into 100□µm sections using a vibratome and stored in 1x phosphate buffered saline (PBS) and 0.01% sodium azide. Sections were rinsed in distilled water, mounted on glass slides, briefly dried, and coverslipped with Aquamount (Polysciences, Warrington, PA). Images were acquired using Leica Stellaris 8 at 2048×2048 format using 63x HC PL APO CS2 (NA□=□1.3) oil immersion objective. For every animal, 20 apical and 20 basal branch segments on tertiary branches were imaged. Spine head diameter and branch length were measured using ImageJ. This was done by going through the z-stack and using the line tool to draw the line through the largest part of each spine to obtain the head diameter. The spine head was measured perpendicular to the spine neck, approximately in the middle of the spine head (57).

For the STED imaging experiment, after the last training session, mice were anesthetized with Euthasol (pentobarbital sodium and phenytoin sodium) and transcardially perfused with warm 0.1□M phosphate buffer, followed by 9% glyoxal (Sigma Aldrich, St. Louis, MO) as described in the following protocol (58). Brains were postfixed in glyoxal overnight at 4□°C. Brains were sectioned into 10□µm cryosections using a cryostat and directly mounted onto glass slides. For immunohistochemistry, sections were rinsed in PBS, non-specific binding was blocked with 3% normal horse serum (heat inactivated, ThermoFisher, Waltham, MA) and 0.3% Triton-X 100 (Sigma-Aldrich) in PBS (sterile filtered). Antibodies/nanobodies were incubated overnight at 4□°C or for 90 minutes at room temperature. GFP (anti-rabbit) was used at 1:250 dilution (Synaptic Systems, Göttingen, Germany) and detected using donkey anti-rabbit Alexa 488 (ThermoFisher, Waltham, MA). Sections were also stained with FluoTag®-X2 anti-PSD95 AZDye568 at 1:500 dilution (Synaptic Systems, Göttingen, Germany). A Leica Stellaris 8 3D tau-STED confocal and super-resolution system (Leica Microsystem) equipped with a tunable white light laser (WLL), CW 592□nm, CW 660□nm, and pulsed 775□nm STED depletion lines were used for image acquisition. HyD-X detectors set to photon counting mode at a 12 mV threshold were used to capture single photons for STED image acquisition. The 100× oil immersion objective (1.4 NA) with 2.7× digital zoom to obtain ∼20□nm pixel size was used to acquire image stacks at 50□nm intervals using 8000 Hz scanning of 2048 × 2048-pixel format images (∼40 µm × 40 µm). 16 line accumulations were used to image individual channels using a resonant (8000 Hz) scanner. Z-stacks for each channel were acquired at 150 nm intervals using sequential imaging with the “between stacks acquisition” setting. Target proteins labeled with nanobodies or antibodies were acquired using the Fast Lifetime Contrast (FALCON) enabled tau-STED module (Leica) with the time-gate on HyD-X detectors adjusted between 0.1 and 6 nanoseconds. GFP labeled proteins were excited using the 491□nm laser (10% maximal AOBS power). The CW 592□nm line (20% AOBS) was used to generate STED. PSD95 labeled with AZDye568 was excited using 579 nm laser (10% AOBS) and the pulsed 775 nm STED laser (15% ABOS) was used to generate STED. All data shown were imaged using 3D STED with 20% STED laser power re-directed toward the Z-donut. For dual color STED imaging, PSD-95 channel (AZDye 568) was acquired first followed by the acquisition of the GFP channel (Alexa 488). Following image acquisition, background photons were removed from raw STED images by adjusting tau strength using the tau-STED module in the FALCON application on the Stellaris 8 STED instrument (Leica Microsystems) as described in detail in (35). Briefly, tau strength was adjusted to the value of 100 to reduce blur surrounding cluster edges in a manner that did not distort existing clusters or create cluster artifacts and it was kept constant throughout all experiments. Afterward, tau-STED images were adjusted to generate high-contrast images for figures and analysis. Tau-STED images were obtained as 16-bit fiff files. These images were initially adjusted in ImageJ by subtracting the background (mean pixel intensities of the entire 2048 × 2048 frame) from every channel. Individual ROIs for every spine were drawn, and experimenters counted every PSD95 puncta that was at least 50% overlapping with the spine, blinded to the experimental condition. Individual PSD-95 clusters were determined an unbiased manner by performing automated 3D segmentation using a custom built macros in image J. Briefly, PSD-95 puncta were considered a separate entity if the line profile between two neighboring clusters was larger than mean + 1.5 standard deviations of fluorescence intensities of all pixels within the spine ROI.

For *in vivo* imaging, mice were placed on a head pad (37□°C) and kept lightly anaesthetized with 0.5% isoflurane. Imaging was performed using a custom-built two-photon microscope equipped with a resonant galvo scanning module (Thorlabs), controlled by ScanImage (http://scanimage.org). The light source was a Mai Tai Deep See ultra-fast Ti-Saph tunable laser (690-1040 nm) (Spectra-Physics) running at 920□nm. The objective was a 20x long distance 1.0 NA water immersion objective (Olympus). Volumetric imaging was done with a piezo (Physik Instrumente). The power used was 40–50□mW. For GFP imaging, z-stacks of branches were taken with 0.5 µm step size with 1024×1024 pixel resolution. For gCaMP imaging, single plane images were collected at 15□Hz, with 1024×1024 pixel resolution. The orientation, curvature and the branching pattern of the dendrites together with the constellation of spines, helped to precisely identify the same field of view in long-term imaging experiments.

For analysis of *in vivo* GFP imaging, Osirix software (https://www.osirix-viewer.com/) was used to line up images from the same dendrites at different timepoints (59). Spines that were present in every time point were identified, then new and eliminated spines were identified. The fraction of stable spines was quantified by taking the number of spines in a timepoint and dividing them by the number of stable spines in the previous timepoint. This was done to quantify the percentage of newly formed and eliminated spines as well.

For analysis of the gCaMP expressing spines, image stacks were motion corrected using the available module in the AUTOTUNE software (60). Regions of interest (ROI) were then drawn around every spine, dendritic branch, or neuron, and the mean gray value for each frame was calculated for every ROI. ΔF/F_0_ was calculated for each ROI. First, the F_0_ was calculated by averaging the fluorescence of the spine/dendrite/neuron while the animal was viewing a blank screen. Next, for every frame, the F_0_ was subtracted, then divided from the mean gray value of that frame to get the ΔF/F_0_. For spines, to ensure that potentially contaminative events from back-propagating action potentials were excluded, we used a subtraction procedure as described in previously published work (13,17,19,61). After extracting the ΔF/F_0_ of the spine and parent dendrite, we plotted ΔF/F_0_^spine^ versus ΔF/F_0_^dendrite^. We fitted a line using linear regression and multiplied the slope of this line by ΔF/F_0_^dendrite^. We then subtracted the scaled dendritic signal from the spine signal to get a spine specific signal. The spine-specific signals were then aligned with the timestamps for visual stimuli. Next, a threshold was set for each spine by multiplying the S.D. of the ΔF/F_0_^spine^ by 3, and threshold crossings were marked. For orientation selectivity, custom-written scripts in Spike2 were used to plot the baseline-subtracted responses of spines to the presented orientations and construct a tuning curve, which was then fitted with a Gaussian to determine the preferred (O_pref_) and orthogonal (O_orth_) orientations, and calculate the orientation selectivity index (OSI) as (R_pref_-R_orth_)/(R_pref_+R_orth_), where R is the event rate at preferred and orthogonal orientations. Tuning width was calculated as half-width at half-maximum of the O_pref_. We considered spines orientation selective if their peak activity was 180° apart, and their OSI was > 0.3. The baseline-subtracted responses to each orientation were plotted as response magnitude. Vector sum calculation for orientation selectivity was done as described previously (40)

### Quantification and statistical analysis

All analyses were performed with the researchers blind to the condition. Statistical analyses were performed in Spike2, ImageJ, Microsoft Excel, MATLAB, JASP(Version0.97.0 Windows, JASP Team, 2026), and GraphPad Prism 9.0 (GraphPad Inc., La Jolla, USA) using as indicated in text and figure legends. All data are reported as mean ± SD, where N represents the number of animals, dendritic branches, or spines, as indicated. Target power for all sample sizes was 0.8 and alpha was set to 0.05.

## Acknowledgements

This study was supported by the Brain Institute Presidential Fellowship to M.F., the Department of Psychology at the University of Virginia start-up funds, the National Center For Advancing Translational Sciences of the National Institutes of Health under Award Numbers KL2TR003016/ULTR003015, Owens Family Foundation and R01MH140184 to A.R. L.Y. D. and D.G.E. were in part supported by Summer Research Internship Program (SRIP). The authors would like to thank the Program in Fundamental Neuroscience for access to the Leica Stellaris confocal microscope.

