## Supplementary Information for "Robust learning-driven structural and functional plasticity of spines in the mature mouse cortex"

**Supporting Information**

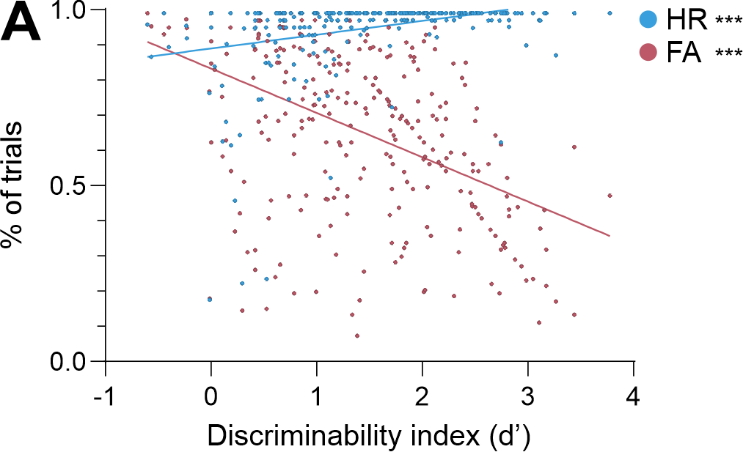

**S1 Fig: Improved task performance is based on an increase in the percentage of hit rate, and decrease in the percentage of false alarm trials. A.** Correlation between discriminability index (d’) versus hit rate (blue) or false alarm (red) for every training session (Pearson’s correlation, hit trials r=0.314, p=8.751x10^-8^_,_ false alarm trials r=-0.458, p=7.253x10^-16,^ N=22 mice).

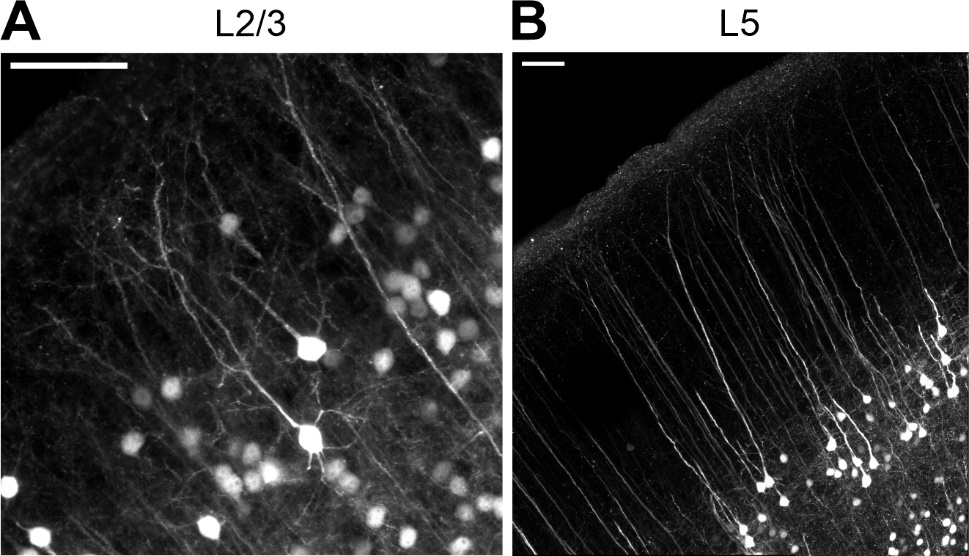

**S2 Fig: Layer 2/3 and Layer 5 Representative Neuron Images. A.** Representative image showing viral labeling of a layer 2/3 neuron in the V1. Scale bar = 50 micrometers. **B**. Representative image showing eYFP in layer 5 neurons in V1. Scale bar=50 micrometers.

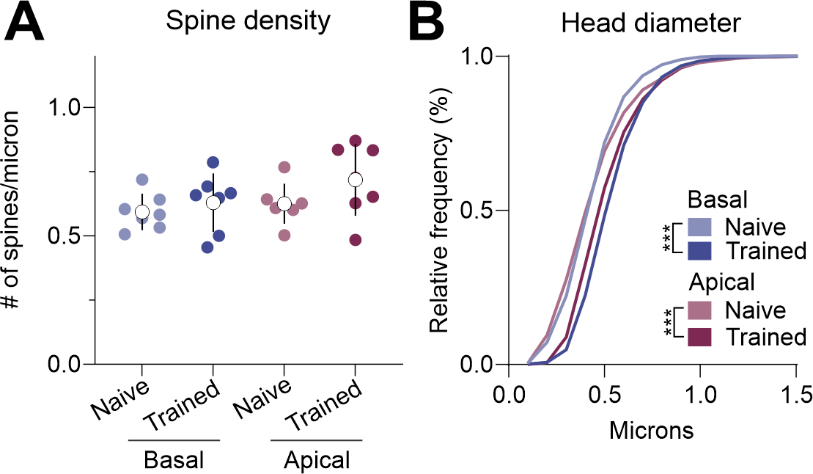

**S3 Fig: Training results in increased spine density and spine head size of layer 5 neuron spines.** **A.** Spine density quantification for naive and trained group for basal (blue) versus apical (magenta) branches (LMM, Training effect p=0.047, AIC=-55.751, BIC=-24.892, Post hoc Apical Naive vs Trained p=0.127, Basal Naive vs Trained p=0.127, N=7 naive, 7 trained mice). **B.** Cumulative frequency histograms of head diameters for basal (blue) and apical (magenta) branches (Kolmogorov-Smirnov test, Apical p=2.2x10^-16^, D=0.19603, Basal p=2.2x10^-16^, D=0.25847, N=7 naive, 7 trained mice).

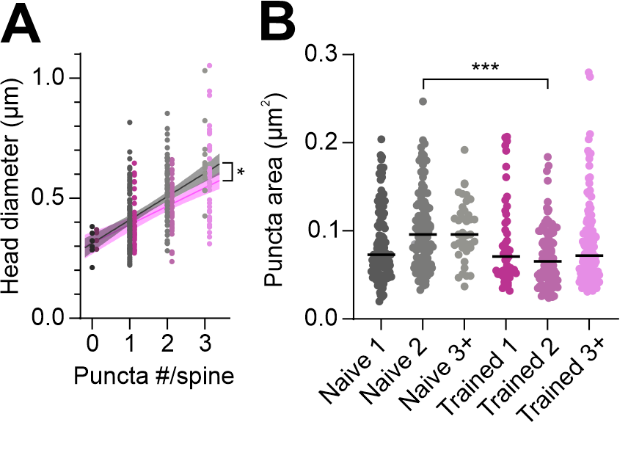

**S4 Fig: STED imaging reveals a positive correlation between head diameter and number of PSD puncta per spine, with spine with multiple PSDs having smaller PSD area. A.** Correlation between spine head size and number of PSD95 puncta for the naive (gray) or trained (magenta) group (Pearson’s correlations, Naive Pearson’s r= 0.5516, p=5.55x10^-20^, Trained Pearson’s r= 0.5348, p=3.350x10^-16^,N=187 spines naive, 139 spines trained, 3 mice per group). **B.** Difference in puncta area between naive (gray) or trained (magenta) groups for spines with 1,2, or 3+ puncta (LMM, Training effect p=0.408, Puncta # effect p=0.99, Training*Puncta # effect p=0.047, AIC=-1826.03, BIC=-1770.482, Naive 2 puncta spine vs Trained 2 puncta spine P_holm_=6.876x10^-4^ N=187 spines from 3 naive mice, 139 spines from 3 trained mice).

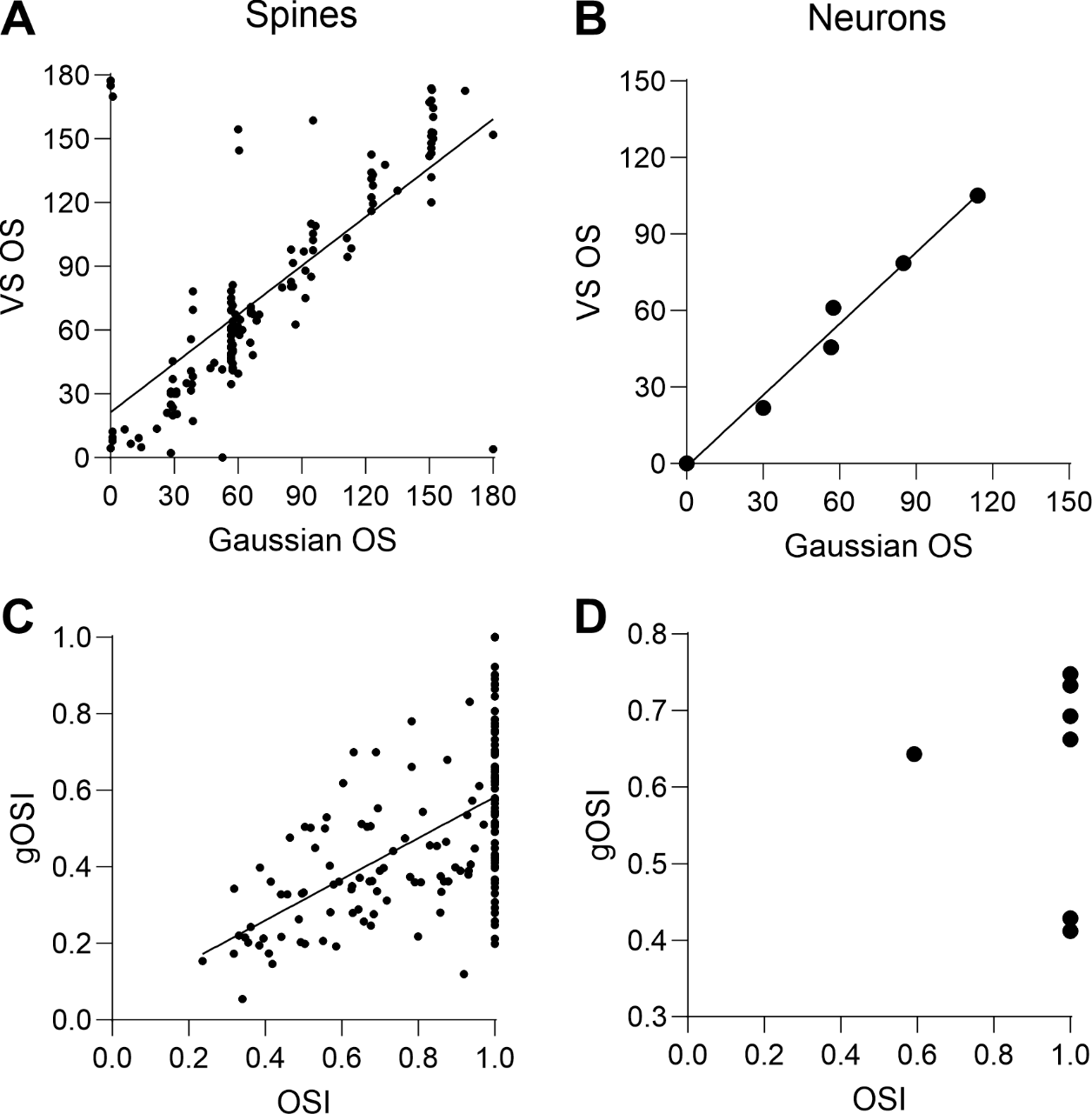

**S5 Fig: Vector sum versus gaussian calculation for orientation selectivity. A.** Orientation selectivity preference calculated in two different ways (gaussian versus vector sum) for the same spines, and neurons (**B**). OSI calculated via vector sum or gaussian OS for spines (**C**) and neurons (**D**). (A, Pearson’s correlation, r=0.715, p=1.114x10^-25^, N=156 spines, B, Pearson’s correlation, r=0.993, p=8.765x10^-6^, N=7 neurons, C, Pearson’s correlation, r=0.573, p=1.039x10^-15^, N=156 spines, D, Pearson’s correlation, r=-0.082, p=0.861, N=7 neurons).

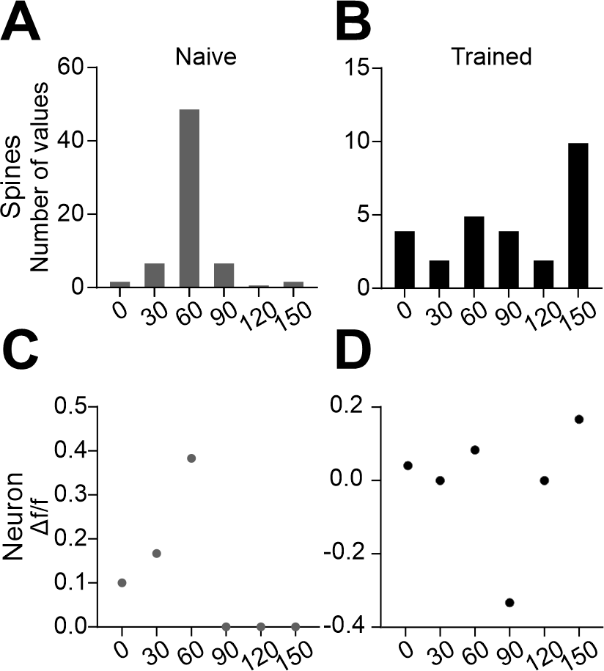

**S6 Fig: Distribution of orientation selection spines reflects the orientation selectivity preference of the host neuron for naive and trained groups. A.** The distribution of orientation selectivity preferences of spines from the same host neuron during the Naive and **(B)** Trained imaging time point. **C.** The activity of the host neuron to each orientation during the Naive, and **(D)** Trained imaging timepoint.

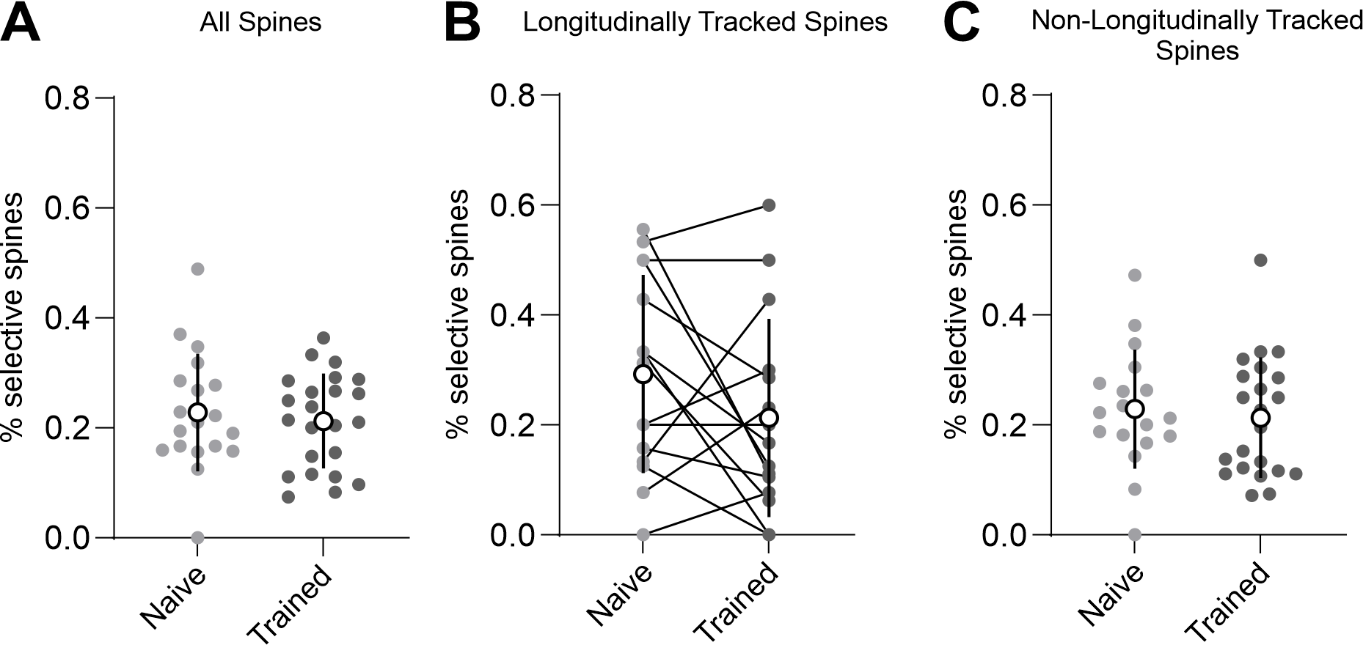

**S7 Fig: Learning does not result in changes in the overall percentage of selective spines across the naive and trained timepoints.** The percentage of selective spines per branch in the naive (light gray) versus trained (dark gray) imaging session for all spines (**A**), longitudinally tracked spines (**B**), or non-longitudinally tracked spines (**C**). (A. LMM, p=0.602, AIC=-58.413, BIC=-51.462 N=23 branches, B. Repeated measures ANOVA, F^(1,14)^= F_training_=2.179, p=0.162, N=13 branches, C. LMM, p=0.645, AIC=-45.233, BIC=-38.477 N=23 branches).

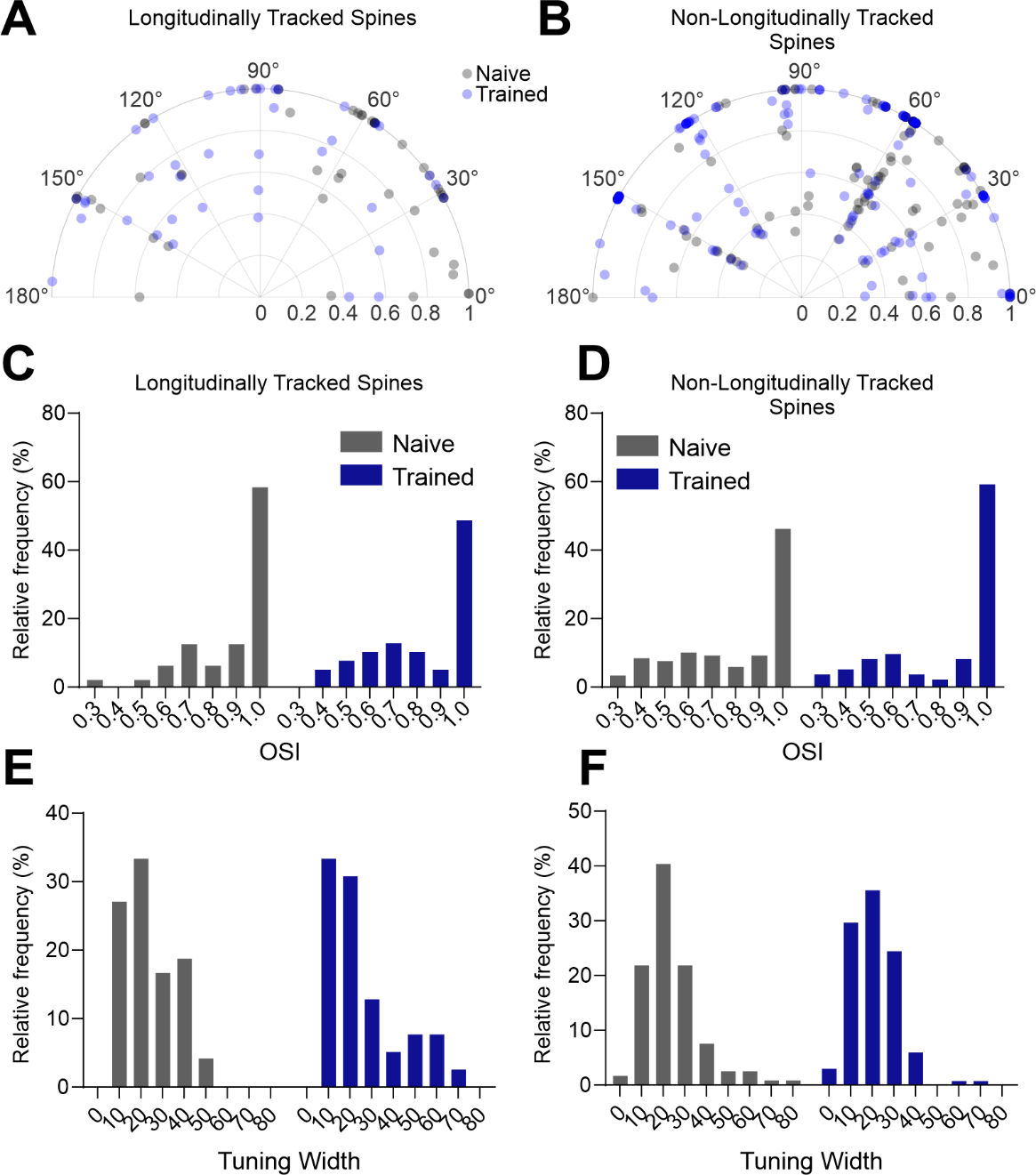

**S8 Fig: Learning does not change orientation selectivity index or tuning width for longitudinally and non-longitudinally tracked spines. A.** Polar plot showing the orientation selectivity preference as well as orientation selectivity index (OSI) during naive (grey) and trained (blue) timepoints for longitudinally and **(B)** non-longitudinally tracked spines. **C.** OSI distribution of spines during naive (grey) and trained (blue) timepoints for longitudinally and **(D)** non-longitudinally tracked spines. (Kolmogorov-Smirnov Test, (C) longitudinally tracked spines, p=0.1739, D=0.20673, (D) non-longitudinally tracked spines, p=0.09933, D=0.15406, (E) longitudinally tracked spines, p=0.4306, D=0.17949 (F) non-longitudinally tracked spines, p=0.182, D=0.13763, longitudinally tracked spines N=40 spines naive, 49 spines trained, non-longitudinally tracked spines N=119 spines naive, 135 spines trained).

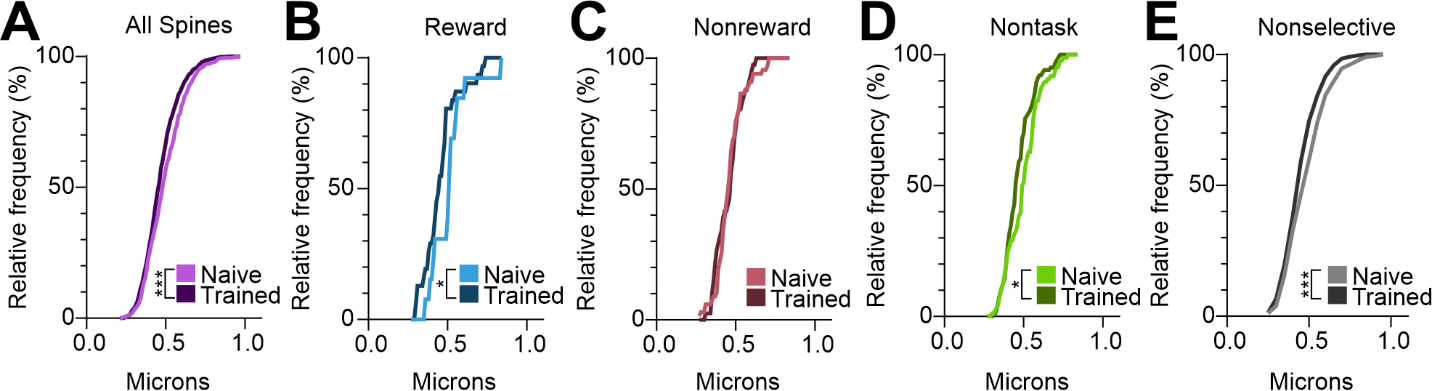

**S9 Fig: Learning results in reduced head diameter of gCaMP labeled neurons, specifically for spines tuned to the reward and nontask orientations. A.** Distribution of head diameters for all gCaMP labeled spines, regardless of tuning (Kolmogorov-Smirnov Test, p=4.223x10-5, D=0.12153, N=648 spines naive, 833 spines trained). **B.** Distribution of head diameter for spines tuned to the rewarded orientation in naive and trained imaging session (Kolmogorov-Smirnov Test, p=0.01288, D=0.49876, N=13 spines naive, 31 spines trained). **C.** Distribution of head diameter for spines tuned to the nonrewarded orientation in naive and trained imaging session (Kolmogorov-Smirnov Test, p=0.4304, D=0.16382, N=67 spines naive, 41 spines trained). **D.** Distribution of head diameter for spines tuned to the nontask orientations in naive and trained imaging session (Kolmogorov-Smirnov Test, p=0.0115, D=0.22782, N=86 spines naive, 103 spines trained). **E.** Distribution of head diameter for spines that are nonselective in naive and trained imaging session (Kolmogorov-Smirnov Test, p=1.33x10^-5^, D=0.1492, N=451 spines naive, 659 spines trained).

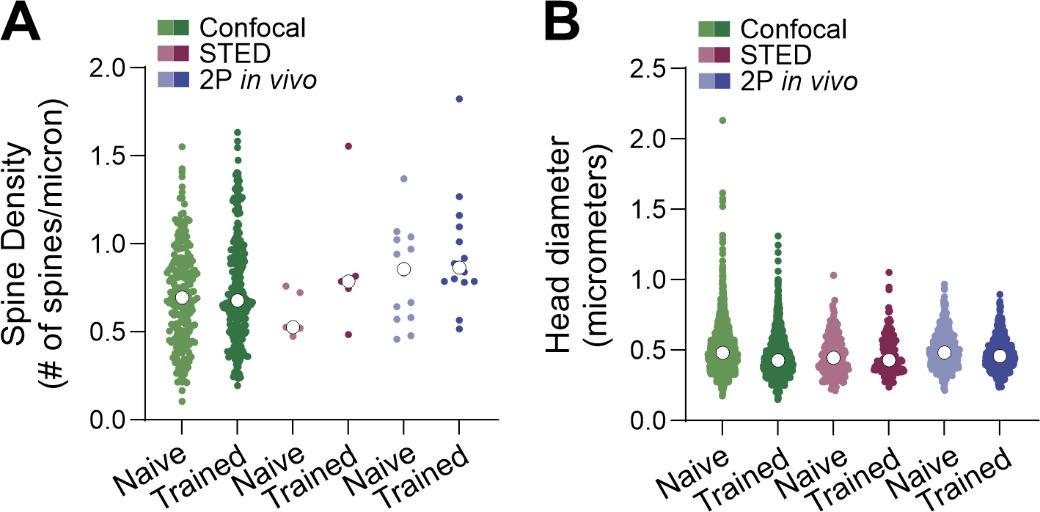

**S10 Fig: Spine density and head diameter are not significantly different for layer 2/3 neurons across different imaging techniques or experiments. A.** Spine density quantification for naive and trained group for layer 2/3 confocal experiment (green), STED experiment (magenta), and 2-photon *in vivo* imaging experiment (blue) (LMM, Training effect p=0.203, Experiment effect p=0.529, AIC=91.193, BIC=125.376, N= 211 branches Naive confocal, N=281 branches Trained confocal, N=5 branches STED, N=14 branches 2P *in vivo*). **B.** Spine head quantification for naive and trained group for layer 2/3 confocal experiment (green), STED experiment (magenta), and 2-photon *in vivo* imaging experiment (blue) (LMM, Training effect p=0.011, Experiment effect p=0.748, AIC=-9054.935, BIC=-8984.296, N= 2958 spines naive confocal, N=3909 spines trained confocal, N=187 spines naive STED, N=139 spines trained STED, N=612 spines naive 2P *in vivo,* N=833 spines trained 2P *in vivo*).

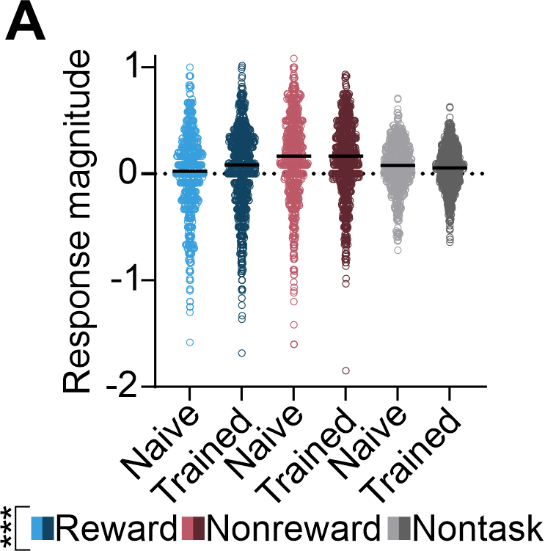

**S11 Fig: Response magnitude is significantly different for different visual stimuli. A.** Response magnitude of spines to the rewarded, non-rewarded, and non-task orientations during the naive and trained imaging session (Generalized linear mixed model, stimulus p=1.307x10^-5,^ AIC=2828.921, BIC=2924.897, N=648 spines Naive, 833 spines Trained).

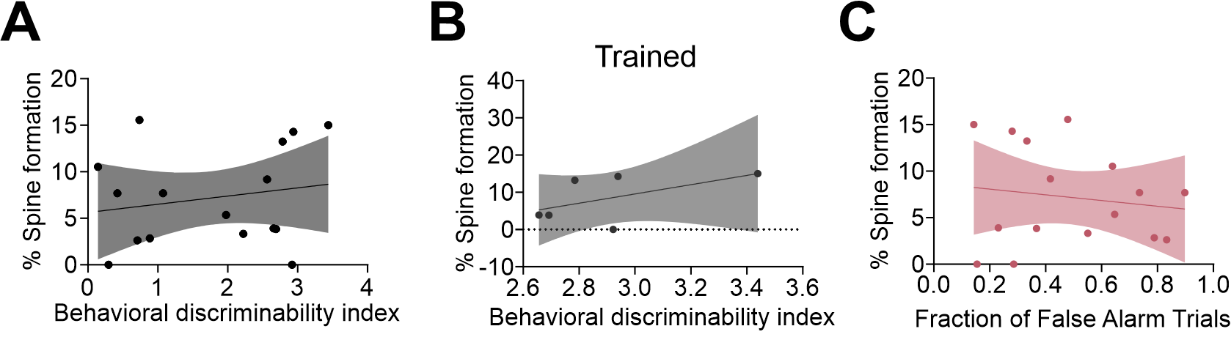

**S12 Fig: Behavior is not correlated with structural changes in spines. A.** Correlation between the behavioral discriminability index for all imaging sessions and the corresponding percentage of new spines formed (Pearson’s correlation, r=0.1886, p=0.4842, 6 animals for day 1 and post time points, N=4 animals for day 5 time points for 16 points overall). **B.** Correlation between behavioral discriminability index for trained timepoint and the percentage of new spines formed (Pearson’s correlation, r=0.5452, p=0.2632, N=6 animals). **C.** Correlation between the fraction of false alarm trials for all imaging sessions and the corresponding percentage of new spines formed (Pearson’s correlation, r=-0.1437, p=0.5955, 6 animals for day 1 and post time points, N=4 animals for day 5 time points for 16 points overall).
